# Alkyne modified purines for assessing activation of *Plasmodium vivax* hypnozoites and growth of pre-erythrocytic and erythrocytic stages in *Plasmodium* spp

**DOI:** 10.1101/2021.10.12.464062

**Authors:** Alona Botnar, Grant Lawrence, Steven P. Maher, Amélie Vantaux, Benoît Witkowski, Justine C. Shiau, Emilio F. Merino, David De Vore, Christian Yang, Cameron Murray, Maria B. Cassera, James W. Leahy, Dennis E. Kyle

## Abstract

Malaria is a major global health problem which predominantly afflicts developing countries. Although many antimalarial therapies are currently available, the protozoan parasite causing this disease, *Plasmodium spp.,* continues to evade eradication efforts. One biological phenomenon hampering eradication efforts is the parasite’s ability to arrest development, transform into a drug-insensitive form, and then resume growth post-therapy. Currently, the mechanisms by which the parasite enters arrested development, or dormancy, and later recrudesces or reactivates to continue development, are unknown and the malaria field lacks techniques to study these elusive mechanisms. Since *Plasmodium spp.* salvage purines for DNA synthesis, we hypothesized that alkyne-containing purine nucleosides could be used to develop a DNA synthesis marker which could be used to investigate mechanisms behind dormancy. Using copper-catalyzed click chemistry methods, we observe incorporation of alkyne modified adenosine, inosine, and hypoxanthine in actively replicating asexual blood stages of *P. falciparum* and incorporation of modified adenosine in actively replicating liver stage schizonts of *P. vivax.* Notably, these modified purines were not incorporated in dormant liver stage hypnozoites, suggesting this marker could be used as a tool to differentiate replicating and non-replicating liver forms and, more broadly, a tool for advancing our understanding *Plasmodium* dormancy mechanisms.

## 1. INTRODUCTION

Malaria is caused by parasites of the genus *Plasmodium* and remains a major global health problem that infects 218 million and kills about 409,000 people a year, mostly children under the age of five. While much effort has been made over the years towards the control and elimination of this disease, recent progress has plateaued [1]. Of the five species affecting humans, *P. falciparum* and *P. vivax* account for most cases and pose the greatest threat. Though *P. falciparum* is the deadliest of the human infecting species, *P. vivax* is more widespread geographically and produces a dormant uninucleate liver stage. Termed hypnozoites, these forms can persist for days, months, and even years, before an unknown mechanism stimulates their reactivation and causes relapsing infections [2]. *P. falciparum* does not produce hypnozoites, however, a stress-induced growth arrest in the asexual blood stage ring stage parasites has been observed when treated with artemisinin (ART) monotherapy [3–5]. This mechanism of induced quiescence is theorized to be a means for drug evasion and subsequent parasite recrudescence once drug pressure is removed. Both species present their own unique obstacles, in terms of eradicating malaria, with quiescence being a common thread.

Hypnozoites are insensitive to all currently marketed antimalarials except the 8-aminoquinolones primaquine and tafenoquine. However, these drugs are contraindicated in patients with a glucose-6-phosphate dehydrogenase (G6PD) deficiency or who are pregnant [6, 7]. This highlights the urgency of discovering novel anti-malarial drugs, the development of which would be aided by a better understanding of the hypnozoite’s basic biology. Whilst suitable high-throughput *in vitro* assays for screening compounds against *P. vivax* liver stage parasites have been recently developed, to date there are no specific markers to exclusively distinguish hypnozoites from liver schizonts [8]. A recombinant antibody reactive to the parasitophorous vacuole membrane (PVM) resident protein upregulated in Infectious Sporozoites 4 (PvUIS4) has been generated, but it immunofluorescently stains both hypnozoites and liver schizonts [9]. Differentiation between the two forms relies on parasite size and specific chemosensitivity, thus much care must be taken to morphologically distinguish hypnozoites from early schizonts [10]. In addition to hypnozoite-specific markers, the identification of markers indicating reactivation of DNA synthesis in hypnozoites would aid in characterizing the mechanism of dormancy and resumption of growth. Two examples of similar markers are Liver-Specific Protein 2 (LISP2), and acetylated lysine 9 of histone H3 (H3K9Ac) [11, 12]. LISP2 has been found to express in early developing liver stage parasites; however, it is limited in that staining is not observed until three days post-infection in *P. vivax.* Furthermore, while LISP2 expression is an early event in liver stage schizont development, the timing of increased expression of LISP2 versus DNA replication has not yet been characterized [13]. H3K9Ac has been elegantly used to accurately count individual nuclei in hypnozoites versus liver stage schizonts [12]. However, this marker indicates only nuclear division and not necessarily active DNA synthesis. In order to differentiate and capture hypnozoites at the moment of reactivation, a marker for DNA synthesis is needed.

Acute, uncomplicated *P. falciparum* infections are most often treated with ART combination therapies (ACT) that are active against blood stages [14]. ART derivatives are fast acting drugs and are extremely effective in reducing parasite biomass. While this treatment has been extremely effective in reducing malaria burden, slower parasite clearance times have been reported as resistance to ART treatment begins to rise [15–18]. Failure rates of ART monotherapy vary widely, anywhere from 2%-50%, and these have been shown to associate with developmentally arrested ring stages [19–21]. While *in vitro* culturing of *P. falciparum* asexual blood stage and induction of dormancy is possible, elucidating the underlying mechanisms of the induced dormant rings remains technically challenging. These dormant parasites present phenotypically with condescend nuclei and reduced cytoplasm, and thus are difficult to differentiate from dead parasites. Although much work has been done to provide insights into DHA-induced dormancy, the molecular mechanism that allows some parasites to enter this stage and later recrudesce is unknown. Furthermore, these dormant stages recrudesce asynchronously, and we currently lack the tools to differentiate between early versus late activators. The development of a DNA synthesis marker that differentiates latent versus active parasites would aid in studying how DHA-induced dormant parasites recover from quiescence.

In many organisms, 5-bromo-2’-deoxyuridine (BrdU) and ethynyl-2’-deoxyuridine (EdU), analogs of the nucleoside thymidine, have been used to identify proliferating cells versus non-proliferating cells [22, 23]. Previous attempts have been made to adapt these labelling techniques for *Plasmodium,* however they failed due to the parasite’s requirement for de novo synthesis of pyrimidines [24]. Although *P. falciparum* expresses transporters that should allow BrdU to be taken up [25, 26], the parasite lacks a thymidine kinase (TK) and thus cannot convert thymidine from a deoxynucleoside into a deoxynucleotide. Recently, studies showed that transfection with TK can allow BrdU labelling in *P. falciparum,* however parasites became much more sensitive to BrdU toxicity [27]. Furthermore, this technique is not suitable for *Plasmodium* species that cannot be easily cultured for transfection, such as *P. vivax. Plasmodium* is a purine auxotroph however, and thus salvages host cell purines [28]. Therefore, similar labeling techniques using alkyne modified purines instead of pyrimidines should be amenable to study *Plasmodium* biology. Mammalian cells incorporate alkyne modified purine versions of adenosine, 7-deaza-7-ethynyl-2’-deoxyadenosine (EdA) and guanosine, 7-deaza-2’-deoxyguanosine (EdG), and recent work with a related apicomplexan, *Cryptosporidium parvum,* showed incorporation of EdA [29, 30]. Thus, we hypothesized alkyne modified purines present a potential DNA synthesis marker that can be designed to differentiate between active versus proliferating and dormant versus nonviable parasites. The sequential steps of *P. falciparum’s* metabolism of adenosine to inosine via an adenosine kinase (AK) followed by the conversion from inosine to hypoxanthine via the purine nucleoside phosphorylase (PNP) presents an additional benefit to synthesize and investigate incorporation of alkyne labeled inosine and hypoxanthine. Hypoxanthine is the closest precursor in the parasite’s metabolic pathway for all purine nucleotides and deoxynucleotides which are used for nucleic acid synthesis.

We hypothesized a modified purine could serve as a marker for reactivation from dormancy as an indicator of DNA synthesis and we designed several purine analogs to be labelled using fluorescent chemo-labelling (“click chemistry”). Click chemistry provides the advantage of no animal reactivity, no potential for cross-reactivity, easier production and storage, and increased flexibility in multicolor experiments. In this study, the development and application of clickable nucleoside analogs EdA, 7-deaza-7-ethynyl-2’-deoxyinosine (EdI), 7-deaza-7-ethynylhypoxanthine (EdH), and 8-ethynylhypoxanthine (8eH), collectively termed EdX, as DNA synthesis markers of proliferating parasites is described. Furthermore, we use EdA staining to help differentiate between dormant and developing liver stage parasites.

## 2. MATERIALS AND METHODS

### 2.1 Chemistry

Given the ability of *P. falciparum* to salvage purines, we hypothesized that alkyne modified derivatives of purine precursors could be developed as tools for the study of DNA synthesis in blood and liver stages of the life cycle. Previous studies with mammalian cells demonstrated that EdA and EdG can be used for cell proliferation studies. For our studies with malaria parasites, we obtained EdA from a commercial source (Carbosynth, United Kingdom), yet similar derivatives of inosine and hypoxanthine required novel synthetic methods (described below).

#### 2.1.1 Synthesis of EdI (3)

##### 7-(2-Deoxy-b-D-erythro-pentofuranosyl)-5-iodo-4-methoxy-7*H*-pyrrolo[2,3-*d*]pyrimidine (2)

To a suspension of 6-chloro-7-deazapurine (**1**, 5.0 g, 33 mmol) in dichloromethane (280 mL) was added *N*-iodosuccinimmide (8.5 g, 38 mmol) and the resulting mixture was stirred at room temperature for 6 hours. The suspension was filtered and concentrated under reduced pressure, and the residue was recrystallized from MeOH to give 6-chloro-7-iodo-7-deazapurine as an off-white solid (5.77 g, 20.7 mmol; 63%). A solution of this product (4.6 g, 16 mmol) was created in acetonitrile (110 mL), and potassium hydroxide (2.6 g, 46 mmol), and tris[2-(2-methoxyethoxy)-ethyl]amine (0.53 mL, 1.6 mmol) were added. After stirring at room temperature for 15 minutes, Hoffer’s chlorosugar (6.8 g, 18 mmol) was added, and stirring was continued for an additional 15 minutes. All insoluble materials were filtered off, and the filtrate was evaporated and suspended in 0.5M sodium methoxide in methanol (55 mL). The suspension was stirred overnight, evaporated and the residue purified by flash chromatography (dichloromethane/methanol 9:1) to give **2** as a colorless solid (4.06 g, 10.4 mmol; 63%).

##### 7-Deaza-7-ethynyl-2’-deoxyinosine (3, EdI)

A solution of **2** (0.245 g, 0.626 mmol) in 2M aqueous sodium hydroxide (19 mL) was heated to reflux for 5 hours. After cooling to room temperature, the solution was carefully neutralized with dilute acetic acid to pH 7, yielding a solid that was filtered and washed with water. Recrystallization from acetonitrile gave 7-(2-deoxy-b-D-erythro-pentofuranosyl)-3,7-dihydro-5-iodo-4*H*-pyrrolo[2,3-*d*]pyrimidin-4-one (0.2 g, 0.5 mmol; 85%) as a colorless solid. A suspension of this material (0.247 g, 0.655 mmol) was created in dry, degassed dimethylformamide (10.4 mL), and copper(I) iodide (0.025 g, 0.131 mmol) palladium tetrakistriphenylphosphine (0.076 g, 0.065 mmol), triethylamine (0.192 mL, 1.38 mmol), and trimethylsilylacetylene (0.907 mL, 6.55 mmol) were added. The reaction was purged with argon before stirring at room temperature for 14 h. The solvents and other volatiles were evaporated under reduced pressure, and the residue was suspended in a mixture of methanol (11 mL) and water (11 mL) with sodium carbonate (0.209 g, 1.97 mmol) and stirred at room temperature for 2 hours. The solvent was evaporated under reduced pressure and the residue was purified by flash chromatography (dichloromethane/methanol 9:1) to give **3** (0.110 g, 0.400 mmol; 61%) as a tan solid. ^1^H NMR (400 MHz, DMSO-*d*β) δ 12.10 (br s, 1 H), 7.94 (s, 1 H), 7.71 (s, 1 H), 6.44 (t, *J*=6.88 Hz, 1 H), 5.27 (d, *J*=3.95 Hz, 1 H), 4.96 (t, *J*=5.32 Hz, 1 H), 4.30 – 4.36 (m, 1 H), 3.99 (s, 1 H), 3.82 (q, *J* = 3.77 Hz, 1 H), 3.48 – 3.60 (m, 2 H), 2.41 (ddd, *J*=13.30, 7.75, 5.75 Hz, 1 H), 2.20 (ddd, *J*=13.13, 5.93, 2.86 Hz, 1 H) ppm. ^13^C NMR (151 MHz, DMSO-*d*6) δ 157.8, 147.3, 145.6, 126.4, 108.2, 98.7, 88.1, 83.7, 81.9, 77.7, 71.2, 62.2, 40.6 ppm. HRMS [M+H]^+^ Calcd for C_13_H_14_N_3_O_4_ 276.0984; found 276.0976.

#### 2.1.2 Synthesis of EdH (6)

##### 7-Iodo-7-deazahypoxanthine (5)

To a solution of 7-deazahypoxanthine (**4**, 1.0 g, 7.4 mmol) in dry dimethylformamide (20 mL) was added bis(trimethyisilyl)acetamide (3.3 g, 16 mmol), and the resulting solution was stirred at 40 °C for two hours. The reaction was cooled to room temperature and N-iodosuccinimide (2.0 g, 8.9 mmol) was added in one portion. The reaction mixture was protected from light and stirred at room temperature until completion. The mixture was poured into water (50 mL) with stirring. A mild exotherm was observed, followed by precipitation of the crude product. After stirring for 1-2 hours, the product was collected by filtration, washed with water, dried, and purified by flash chromatography (dichloromethane/methanol 9:1) to give **5** (0.65 g, 2.5 mmol; 67%) as a colorless solid.

##### 7-Deaza-7-ethynyl-2’-deoxyhypoxanthine (6, EdH)

To a suspension of **5** (0.050 g, 0.19 mmol) in dry, degassed dimethylformamide (3 mL) was added copper(I) iodide (70 mg, 0.038 mmol) palladium tetrakistriphenylphosphine (0.022 g, 0.019 mmol), triethylamine (0.056 mL, 0.402 mmol), and trimethylsilylacetylene (0.238 mL, 1.92 mmol). The reaction was purged with argon before stirring at 55 °C for 14 hours after which thin layer chromatographic analysis (dichloromethane/methanol, 10:1) showed total consumption of starting material. The reaction mixture was diluted with dichloromethane (20 mL) and extracted with water (2 x 15 mL). The organic phase was separated, dried, and concentrated and the residue was suspended in methanol (5 mL) and water (5 mL) with sodium carbonate (0.061 g, 0.571 mmol) and stirred at room temperature for 2 hours. The solvent was evaporated under reduced pressure and the residue was purified by flash chromatography (dichloromethane/methanol 19:1) to give **6** (.017 g, .107 mmol; 55.8%) as a colorless solid. ^1^H NMR (600 MHz, DMSO-d6) δ 12.08 (br s, 1 H), 7.86 (s, 1 H), 7.39 (s, 1 H), 3.93 (s, 1 H) ppm. ^13^C NMR (151 MHz, DMSO-d6) δ 158.6, 148.2, 145.1, 126.6, 107.9, 98.2, 81.1, 78.3 ppm. HRMS [M+H]^+^ Calcd for C_8_H_6_N_3_O 160.0511; found 160.0504.

#### 2.1.3 Synthesis of 8eH (9) 8-Iodo-2’,3’,5’-tris-tertbutyldimethylsilylinosine (8)

To a clear solution of tert-butyldimethylsilyl chloride (2.81 g, 18.6 mmol) and imidazole (2.54 g, 37.3 mmol) in dry dimethylformamide (10 mL) was added inosine (**7**) (1.0 g, 3.7 mmol) in one portion. The reaction mixture was stirred at room temperature overnight, after which thin layer chromatographic analysis (chloroform/ethyl acetate, 1:1) showed total conversion of starting material to a single product. The white slurry was then partitioned into a mixture of water (100 mL) and dichloromethane (100 mL). The organic layer was separated, and the aqueous phase was extracted with dichloromethane (3 x 50 mL). The combined organic extracts were dried over magnesium sulfate, filtered, and evaporated under reduced pressure to afford a white solid, which was purified by flash chromatography (hexanes/ethyl acetate 9:1) to give 2’,3’,5’-tris-tertbutyldimethylsilylinosine as a white powder (2.1 g, 3.4 mmol; 92%). To a solution of diisopropylamine (2.3 mL, 16 mmol) in dry, degassed tetrahydrofuran (16 mL) at −78 °C was added dropwise n-butyllithium (11 mL, 18 mmol, 1.6 N in hexane). This mixture was stirred for 10 min at −78 °C, then a solution of 2’,3’,5’-tris-tertbutyldimethylsilylinosine (2.0 g, 3.3 mmol) in tetrahydrofuran (37 mL) was added dropwise at −78 °C, followed by stirring at −78 °C for an additional 2 hours. Iodine (3.32 g, 13.1 mmol) in tetrahydrofuran (12 mL) was added dropwise until a deep-yellow color persisted. It was stirred for an additional 10 minutes, after which thin layer chromatographic analysis showed completion of the reaction (chloroform/ethyl acetate 1: 1). The mixture was then warmed to room temperature and quenched with a pH 4 buffer of 1 N sodium acetate/acetic acid. The aqueous phase was washed with dichloromethane (3 x 30 mL); the combined organic extracts were washed with 0.5 N sodium bicarbonate, dried over magnesium sulfate, filtered, and evaporated under reduced pressure to give a yellow oil, which was purified by flash chromatography (dichloromethane/ethyl acetate, 9:1) to give **8** as a pale-yellow solid (2.04g, 2.77 mmol; 85%).

##### 8-Ethynylhypoxanthine (9)

To a suspension of **8** (0.5 g, 0.7 mmol) in dry, degassed dimethylformamide (11 mL) was added copper(I) iodide (0.0260 g, 0.136 mmol) palladium tetrakistriphenylphosphine (0.078 g, 0.068 mmol), triethylamine (0.199 mL, 1.43 mmol), and triisopropylsilylacetylene (1.24 mL, 6.79 mmol). The reaction was purged with argon before stirring at 55 °C for 14 hours, after which thin layer chromatographic analysis (dichloromethane/methanol, 10:1) showed total consumption of starting material. The reaction mixture was diluted with dichloromethane (50 mL) and extracted with water (2 x 25 mL). The organic phase was separated, dried, and concentrated to give a brown oil which was suspended in MeOH (15 mL) and 1N aqueous hydrochloric acid (15 mL) before being heated to reflux overnight. The resultant solution was neutralized with saturated sodium bicarbonate and extracted with dichloromethane (3 x 50 mL). The organic phase was separated, dried, and concentrated and the residue was dissolved in tetrahydrofuran (15 mL) before slowly adding tetrabutylammonium fluoride (0.355 g, 1.36 mmol). The mixture was stirred overnight and was monitored by thin layer chromatography (dichloromethane/methanol, 10:1). Upon completion, calcium carbonate and Dowex 50WX8-100 resin were added and the mixture stirred for 45 minutes. The solids were filtered over a pad of Celite and the filtrate evaporated to dryness. The residue was purified by flash chromatography (dichloromethane/methanol, 10:1) to give **9** (0.064 g, .400 mmol; 59%) as a colorless solid. ^1^H NMR (600 MHz, DMSO-d6) δ 14.00 (br s, 1 H), 12.18 (br s, 1 H), 7.96 (s, 1 H), 4.52 (s, 1 H) ppm. ^13^C NMR (151 MHz, DMSO-d6) δ 155.5, 146.2, 132.6, 83.6, 75.0, 70.2 ppm. [M+H]^+^ Calcd for C7H5N4O 161.0463; found 161.0455.

### 2.2 Biology

#### 2.2.1 *In vitro* culture of intraerythrocytic *P. falciparum* parasites

*Plasmodium falciparum* clone W2 (Indochina II) was cultured using standard techniques [31]. Briefly, parasites were maintained at 37°C in hypoxic conditions (90% N_2_, 5% CO_2_, 5% O_2_) at a hematocrit of 2% A+ human red blood cells (RBCs; Interstate Blood Bank, Memphis, TN). Parasites were cultured in complete medium consisting of RPMI1640 supplemented with 25 mM HEPES, 0.24% (w/v) sodium bicarbonate and either a) 10% heat-inactivated A+ human plasma (Interstate Blood Bank, Memphis, TN) or b) 1% (w/v) AlbuMAX II (Thermo Fisher) and 320 μM hypoxanthine (Sigma). Parasite development was monitored with light microscopy of Giemsa-stained blood smears.

#### 2.2.2 EdX incorporation and Cu-catalyzed azide-alkyne staining of active intraerythrocytic parasites

Asynchronous *P. falciparum* was split to 2% parasitemia in a 2% hematocrit and supplemented with 10 μM EdA (Carbosynth, United Kingdom), EdI, EdH, and/or 8eH, and incubated for 48 hours at 37 °C in hypoxic conditions. After incubation, parasites were briefly centrifuged, and the supernatant removed. Infected cells were then fixed in a solution containing 4% paraformaldehyde and 0.05% glutaraldehyde for 15 minutes at room temperature, adapted from Balu et. al. 2010 [32]. Following a wash with phosphate buffer saline (PBS, pH 7.4), parasites were prepared for the azide-alkyne click reaction by permeabilization with 0.1% Triton X-100 for 10 minutes, followed by a 1-hour incubation with 3% BSA at room temperature. Cells were then stained with a freshly prepared staining mix containing 2 mM CuSO_4_ (Sigma), 12 μM Alexa Fluor Azide 488 (Thermo Fisher), and 10 mM sodium ascorbate (Sigma) for 1 hour in the dark at room temperature. Parasites were then washed once with PBS and costained with 10 μg/mL Hoechst 33342 (Thermo Fisher) for 15 minutes followed by three washes with PBS. They were then mounted onto poly-L-lysine coated slides and imaged with a Zeiss Axio Observer.Z1/7. The FIJI plugin, JACoP, was used to generate Mander’s coefficients and Pearson’s coefficient, *r*, for co-localization of Hoechst 33342 nuclear staining and EdX incorporation.

#### 2.2.3 Cytotoxicity assays of EdX on intraerythrocytic parasites

Cytotoxicity of the modified purines was assessed by measuring the increase in parasitemia over time. Briefly, asynchronous *P. falciparum* was split to 0.5% parasitemia in a 2% hematocrit and supplemented with 10 μM EdX. Parasites were incubated at 37 °C in hypoxic conditions and allowed to grow for 5 days. On day 3 parasites were split 1:5 to avoid parasite death due to overgrowth.

Samples were taken daily and fixed with 4% paraformaldehyde and 0.05% glutaraldehyde and stored at 4 °C until all samples were collected. Parasites were then stained with 10 μM Hoechst 33342 for 15 minutes followed by three washes with PBS and percent parasitemia was analyzed via flow cytometry using a Beckman Coulter CytoFLEX. Primary gating was performed based on background fluorescence from uninfected red blood cells to obtain parasite-infected red blood cells as an indication of parasitemia. Results were then visualized via a GraphPad Prism 9 plot where statistical analysis was also conducted using a 2-way ANOVA multiple comparisons test.

#### 2.2.4 Human Subjects Consideration

*P. vivax* isolates were collected into a heparin tube (BD) via venipuncture from human volunteers following approval by the Cambodian National Ethics Committee for Health Research (113NHECR). Protocols conformed to the Helsinki Declaration on ethical principles for medical research involving human subjects (version 2002) and informed written consent was obtained from all volunteers or legal guardians.

#### 2.2.5 *In vitro* liver stage *P. vivax* incorporation of EdX

Primary human hepatocytes (PHH) were infected with *P. vivax* sporozoites as previously described [8]. Briefly, *Anopheles dirus* mosquitos were fed a bloodmeal containing *P. vivax* gametocytes and maintained on a 12:12 L:D cycle and 10% sucrose in water. Two days prior to infection, PHHs (lot BGW, BioIVT) were seeded into collagen-coated 384-well plates (Greiner Bio-One) at a density of 18,000 cells per well. Mosquito salivary glands were aseptically dissected on days 16-21 post feeding to obtain *P. vivax* sporozoites. PHH seeded plates were then infected with 5,000-18,000 sporozoites per well. Infected PHHs were then exposed to EdX on days 5, 6, 7, and 8 post infection (dpi) with 2 μM EdX, 10 μM EdX, and DMSO vehicle control. Media was exchanged daily immediately before EdX exposure. At 9 dpi, cultures were fixed with 4% paraformaldehyde in PBS. Fixed cultures were stained with recombinant mouse anti-*P. vivax* upregulated in infectious sporozoites-4 antibody (rPvUIS4) [9] diluted 25,000-fold in staining buffer (0.03% Triton X-100, 1% (w/v) BSA in PBS) overnight at 4 °C in the dark. Following three washes with PBS, cells were then stained with rabbit anti-mouse Alexafluor488 (Abcam) diluted 1:1000 diluted in staining buffer for 1 hour at room temperature in the dark. Cultures were then washed three times with PBS followed by staining with a freshly prepared staining mix containing 2 mM CuSO_4_, 12 μM Alexa Fluor Azide 594 (Thermo Fisher), and 10 mM sodium ascorbate for 1 hour in the dark at room temperature. The cells were then washed three times with PBS and counterstained with 10 μg/mL Hoechst 33342 for 15 minutes at 37 °C. Cultures were washed once with PBS and stored in PBS prior to automated high content imaging with a 20x objective on an ImageXpress confocal microscope (Molecular Devices, San Jose, CA). Liver stage parasites and host cell hepatocytes were quantified using the MetaXpress software version 6.6.1.42 for ImageXpress. Individual images were also obtained with a 100x objective on a DeltaVision II deconvolution microscope (Applied Precision Inc., Currently Leica Microsystems, Buffalo Grove, IL). Analysis was conducted using GraphPad Prism 9 with an ordinary one-way ANOVA multiple comparisons test and unpaired t-test.

#### 2.2.6 HepG2 EdX Staining

HepG2 hepatoma cells were cultured in collagen-coated T75 flasks in minimum essential medium eagle with Earle’s BSS (MEM Eagle EBSS) from Lonza (Walkersville, MD) supplemented with 10% FBS and 4.4 mM sodium pyruvate at 37 °C with 5% CO_2_. Cells were seeded at 5,000 cells per well in a collagen coated 384 well plate (Greiner Bio-One). EdX (2 μM or 10 μM) and 0.1% DMSO vehicle control were added 24 hours post seed and allowed to incorporate for 48 hours at 37 °C and 5% CO_2_. Cells were then fixed with 4% paraformaldehyde for 15 minutes at room temperature. Fixed cells were stained with freshly prepared staining mix containing 2 mM CuSO_4_, 12 μM Alexa Fluor Azide 488, and 10 mM sodium ascorbate in staining buffer (0.03% Triton X-100, 1% (w/v) BSA in PBS) overnight at 4 °C in the dark. Following three washes with PBS, cells were counterstained with 10 μg/mL Hoechst 33342 for 15 minutes at 37 °C. Cultures were washed once with PBS and stored in PBS prior to automated high content imaging with a 4x objective on a Lionheart imaging system (Biotek). Viability was measured by counting cell nuclei using Gen5 software (Biotek) and statistical analysis was conducted using GraphPad Prism 9 with an ordinary one-way ANOVA multiple comparisons test and unpaired t-test.

#### 2.2.7 *In vitro* liver stage *P. berghei* incorporation of EdA

*An. stephensei* mosquitos were fed a *P. berghei* gametocyte-infected bloodmeal and maintained on 12:12 L:D cycle and 10% sucrose in water. One day prior to infection with sporozoites, HepG2 cells were seeded at 17,500 cells per well in a collagen-coated 384 well plate (Greiner Bio-One). Mosquito salivary glands were aseptically dissected on day 20-22 post feeding to obtain *P. berghei* sporozoites.

HepG2 seeded plates were then infected with 6,000 sporozoites per well. Infected and uninfected cultures were then treated 24 hours post infection (hpi) with 2 μM EdA, 10 μM EdA, and 0.1% DMSO vehicle control. Media was exchanged before EdA treatment. At 48 hpi, cultures were fixed with 4% paraformaldehyde in PBS. Fixed cultures were stained with mouse monoclonal antibody 13.3 anti-Glyceraldehyde-3-Phosphate Dehydrogenase (GAPDH) diluted 10,000-fold in staining buffer (0.03% Triton X-100, 1% (w/v) BSA in PBS) overnight at 4°C in the dark [8]. Anti-GAPDH was obtained from The European Malaria Reagent Repository. Following three washes with PBS, cells were then stained with rabbit anti-mouse Alexafluor488 diluted 1:1000 diluted in staining buffer (0.03% Triton X-100, 1% (w/v) BSA in PBS) for 1 hour at room temperature in the dark. Cells were then washed three times with PBS followed by staining with a freshly prepared staining mix containing 2 mM CuSO_4_, 12 μM Alexa Fluor Azide 594, and 10 mM sodium ascorbate for 1 hour in the dark at room temperature. Cells were then washed three times with PBS and counterstained with 10 μg/mL Hoechst 33342 for 15 minutes at 37 °C. Cells were washed once with PBS and stored in PBS prior to imaging at 40x on a Zeiss Axio Observer.Z1/7 (Pleasanton, CA).

#### 2.2.8 *In vitro P. falciparum* EdA, ^3^H-hypoxanthine, and ^3^H-adenosine incorporation in dihydroartemisinin (DHA)-induced dormant asexual blood stage parasites

*P. falciparum* (W2) asexual blood stage parasites were synchronized to ring stages using 5% D-sorbitol and were then split to 2% parasitemia in a 2% hematocrit. Dormancy was then induced 48 hours post-synchronization with 700nM DHA for 6 hours while parasites were incubated at 37°C in hypoxic conditions. Following three washes with RPMI1640, parasites were re-suspended and cultured in complete media containing 10% heat-inactivated A+ human plasma. Parasite recrudescence and development was monitored daily with light microscopy of Giemsa-stained blood smears and media was changed every 48 hours. Aliquots of 100 μL were transferred daily (up to 10 days post dormancy induction) to a 96 well plate where 5 μCi ^3^H-hypoxanthine (Perkin Elmer), 5 μCi ^3^H-adenosine (Perkin Elmer), or 10 μM EdA was supplemented. These aliquots were incubated for 24 hours at 37 °C in hypoxic conditions after which they were either frozen at −80 °C (for tritiated samples) or fixed with 4% paraformaldehyde and 0.05% glutaraldehyde and stored in 4 °C until all samples were collected (for EdA-labeled samples). Uninfected red blood cells (uRBC) and active asynchronous parasites at a starting 2% parasitemia were used as controls. Once all samples were collected, click chemistry was performed on EdA labelled samples as described above and incorporation was measured via flow cytometry on a Beckman Coulter CytoFLEX (Indianapolis, IN). Tritiated samples were harvested and counted on a Perkin Elmer Microbeta^2^ scintillation counter (Waltham, MA). Data was graphed and analyzed using GraphPad Prism 9.

## 3. RESULTS

### 3.1 Synthesis and characterization of EdI, EdH, and 8eH

The synthesis of EdI was initiated from commercially available 6-chloro-7-deazapurine (**1**, Scheme 1). Selective iodination at the 7-position followed by addition of Hoffer’s chlorosugar and concomitant deprotection and S_N_Ar displacement with sodium methoxide afforded **2**. Following displacement of the methoxy group, Sonogashira installation of the clickable handle with TMS-acetylene and liberation of the terminal acetylene yielded EdI (**3**). In a similar manner, 7-deazahypoxanthine (**4**) was iodinated and affixed with the alkyne to give EdH (**6**). From a cost-effective perspective, it was most pragmatic to prepare 8eH (**9**) from inosine (**7**). Silyl protection of the carbohydrate alcohols allowed smooth iodination at the 8-position to give **8**, which could then be converted to **9** via a modified Sonogashira protocol followed by global deprotection.

### 3.2 Incorporation of EdX analogues into asexual blood stages of *P. falciparum*

To evaluate if EdA, EdI, EdH, and 8eH were incorporated into newly synthesized DNA of *Plasmodium,* we conducted labeling studies using a copper-catalyzed azide-alkyne cycloaddition chemical reaction with a fluorescent azide probe. Using an *in vitro* asynchronous asexual blood stage culture of *P. falciparum,* we observed that the addition of 10 μM of each of the alkyne modified purines resulted in nuclear staining after a 48-hour incubation period, equivalent to a full asexual blood stage life cycle (Figure 1). The observed staining co-localized with nuclear Hoechst 33342 staining, which preferentially binds to the A-T regions of DNA, and a Fiji JACoP Pearson’s coefficient for EdA, EdI, EdH, 8eH was calculated to range from 0.991 to 0.997, thus indicating that the modified purines were incorporated into parasite DNA. Manders’ coefficients for the four images shown in Figure 1 were also calculated and ranged from 0.925 to 0.999.

**Figure 1:**
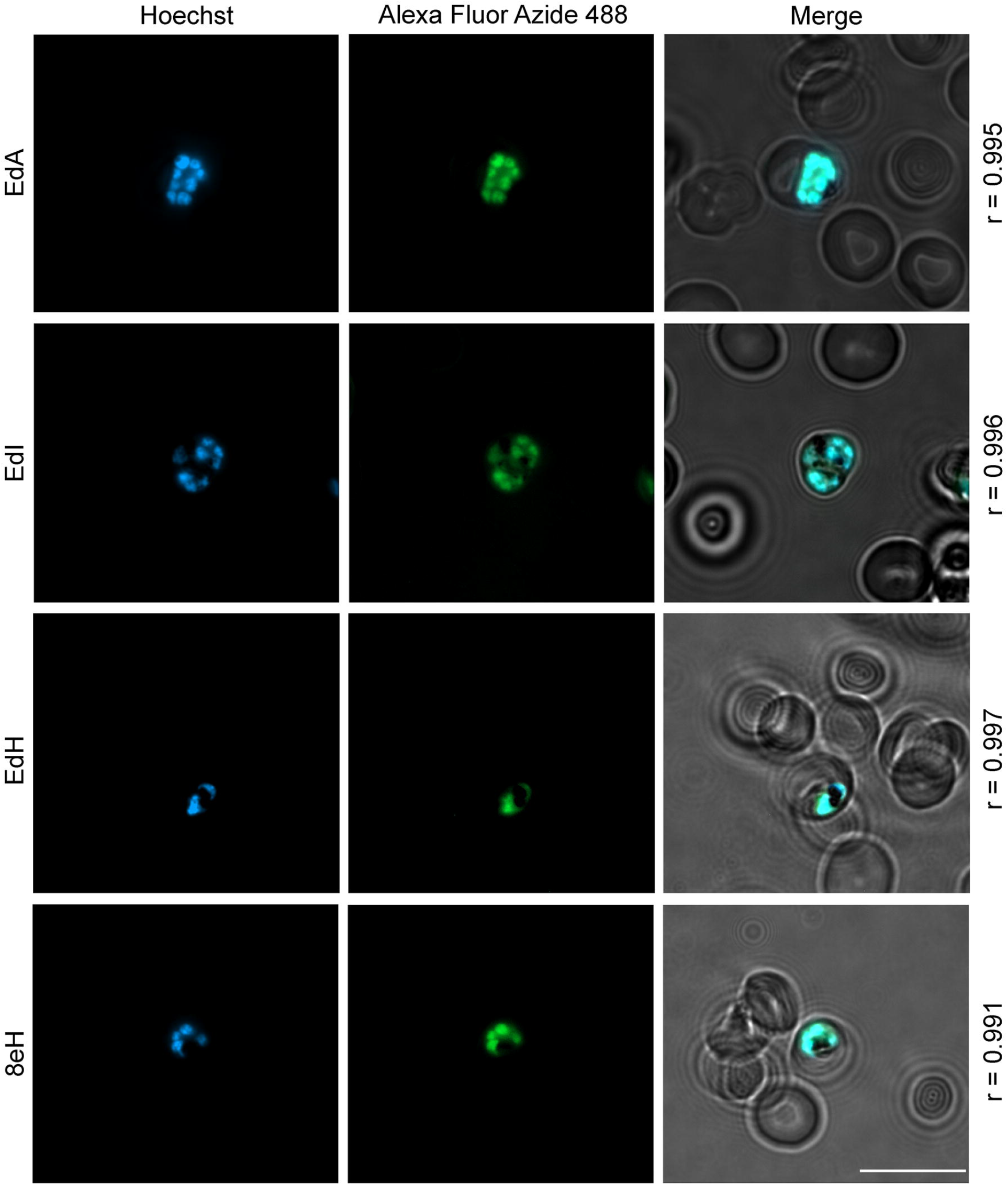
Alkyne modified purines incorporate into the replicating asexual blood stage *P. falciparum.* Detection of modified purines in *P. falciparum* after a 48-hour incubation with 10 μM of EdA, EdI, EdH, and 8eH (green). Parasites were co-stained with 10 μg/mL Hoechst 33342 (blue). Images were obtained on a Zeiss Axio Observer.Z1/7 microscope with a 100x objective. Colocalization was assessed and Pearson’s coefficient (r) was calculated using Fiji JACoP plugin. Scalebar = 10 μm.

Considering the potential cytotoxicity of the modified purines, we assessed parasite growth in the presence of these compounds to quantify their effects on parasite proliferation. Furthermore, because serum supplements contain different levels of purines, we evaluated the cytotoxic effects of EdA, EdI, EdH, and 8eH over a 5-day incubation period in complete media containing albumax II alone (1% w/v), complete media containing albumax II and supplemented with hypoxanthine (10 μM), and complete media containing 5% A+ human plasma. The media containing albumax II alone lacks any purines for the parasites to salvage whereas media supplemented with hypoxanthine or human plasma contains purines, although at different levels. In media without hypoxanthine supplementation, all modified purine compounds showed a deleterious effect (p < 0.0001) on parasite proliferation as compared to the unmodified hypoxanthine media control (Figure 2a). Cultures with media containing either supplemented hypoxanthine or human plasma were unaffected by the addition of alkyne modified purines (Figure 2b-c). These results indicate that while the EdX analogues incorporated into asexual blood stage *P. falciparum* parasite DNA, unmodified purines are necessary for continued parasite growth over time.

**Figure 2:**
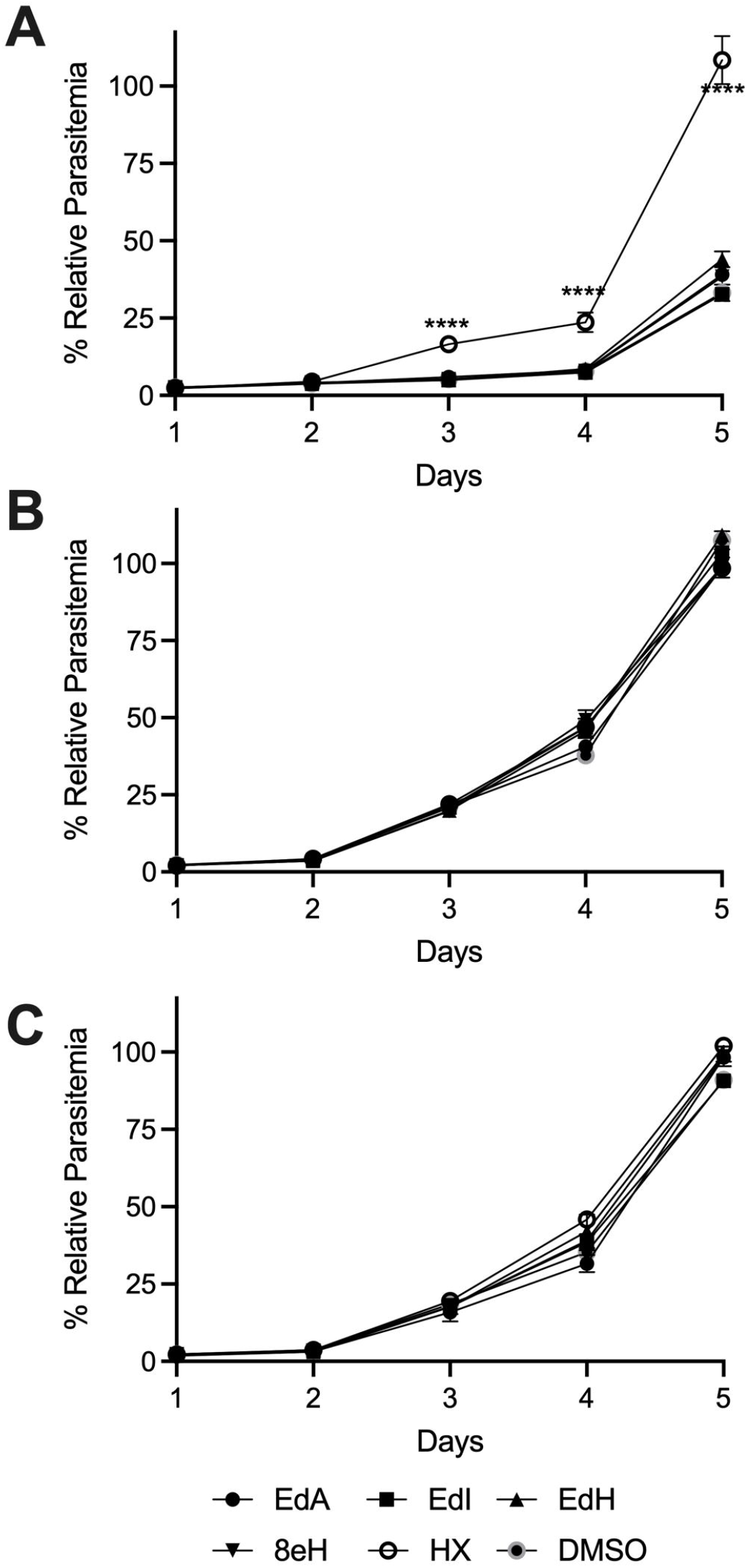
*P. falciparum* asexual blood stage growth depends on unmodified purines in culture media. *P. falciparum* asexual blood stages were grown in media supplemented with 10 μM EdA, EdI, EdH, 8eH, hypoxanthine (HX), or 0.1% DMSO vehicle control in **(A)** complete media supplemented with Albumax II, **(B)** complete media supplemented with Albumax II and hypoxanthine, or **(C)** complete media supplemented with A+ human plasma. Growth was assessed by flow cytometry using a Beckman Coulter CytoFLEX. Data shown are one representative experiment of three independent experiments. Errors (SD) were omitted when smaller than the marker. Significance assessed by 2-way ANOVA and Dunnett’s multiple comparisons test, **** p < 0.0001.

### 3.3 *P. vivax* actively replicating liver stage schizonts incorporate EdA and can be differentiated from dormant hypnozoites via high-throughput content imaging

After invading a hepatocyte, a *P. vivax* sporozoite develops into either an actively replicating liver schizont or a dormant hypnozoite with a single nucleus [33]. Over the first 3-5 days of liver stage culture, hypnozoites and schizonts are of similar sizes and are indistinguishable [10]. Therefore, to verify that EdX analogs incorporate into actively replicating parasite DNA in the liver stage, cultures were supplemented with 2 μM or 10 μM of EdA, EdI, EdH, and 8eH at 5, 6, 7, and 8 dpi and then cultures were fixed at 9 dpi. By using fluorescent microscopy, we observed that the addition of EdA resulted in nuclear staining of only actively replicating liver stage schizonts, whereas dormant hypnozoites did not incorporate EdA. The EdA fluorescence co-localized with nuclear Hoechst 33342 staining, similar to our *P. falciparum* asexual blood stage staining (Figure 3). However, unlike in *P. falciparum* asexual blood stage parasites, *P. vivax* liver stage parasites did not incorporate EdI, EdH, or 8eH (data not shown).

**Figure 3:**
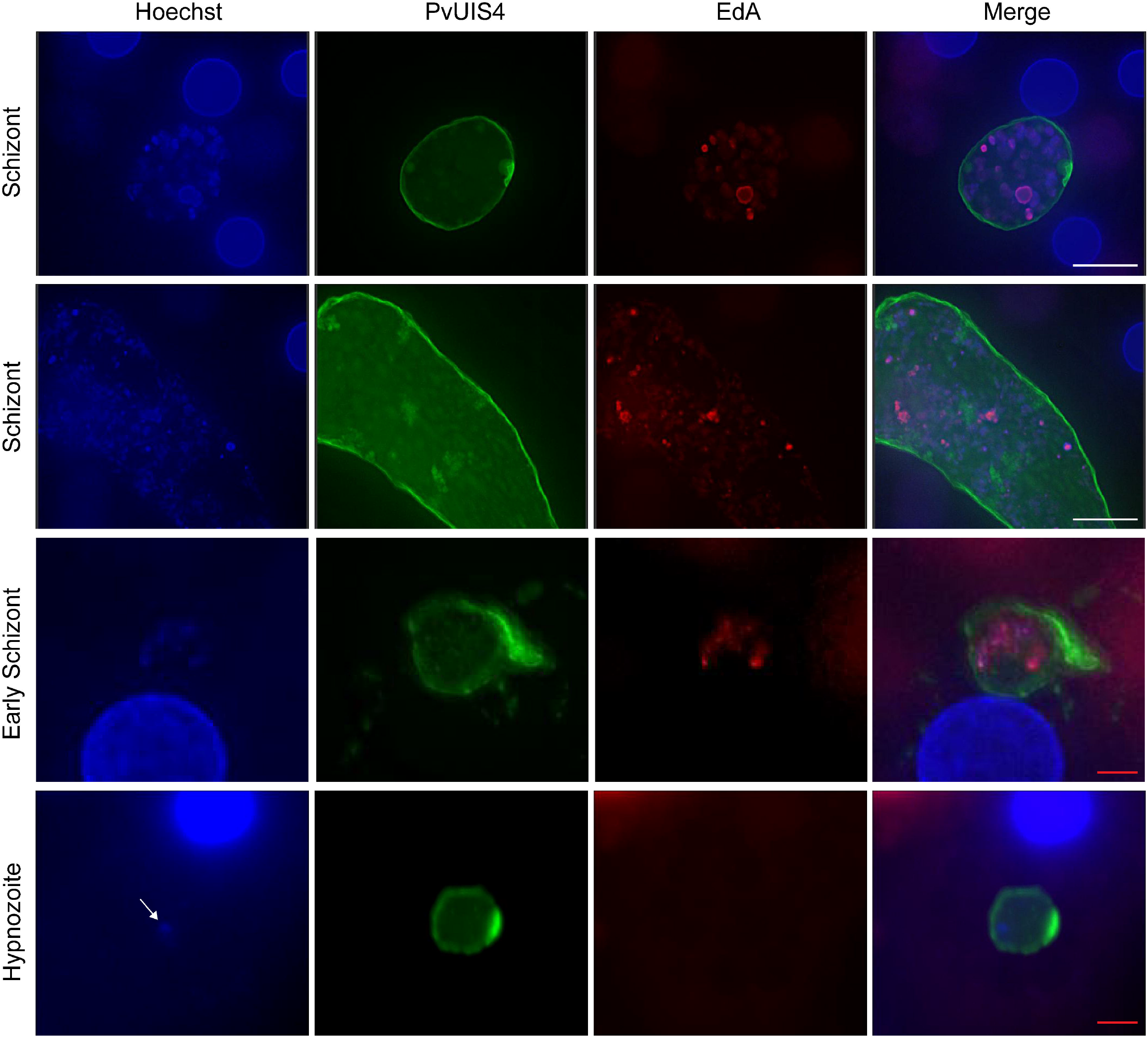
Alkyne modified adenosine (EdA) incorporates in replicating *P. vivax* liver stage parasites, but not in hypnozoites. Primary human hepatocytes were infected with *P. vivax* sporozoites and incubated with 10 μM EdA on days 5-8 post infection. Detection of EdA was assessed via a copper-catalyzed click reaction (red). Parasites were co-stained with 1:25,000 PvUIS4, a parasitophorous vacuole membrane stain (green), and 10 μg/mL Hoechst 33342 (blue). Arrow points to single nucleus in *P. vivax* liver stage hypnozoite. Images were obtained on a DeltaVision II deconvolution microscope at 100x objective. White scalebar = 15 μm. Red scalebar = 5 μm.

We next evaluated if the EdA staining in *P. vivax* liver stage schizonts could be identified in a high-throughput manner using an automated high content imaging system. We tested two concentrations of EdA and observed that separation of liver stage schizonts and dormant hypnozoites can be accomplished using 2 or 10 μM EdA (Figure 4a-b). Since all *P. vivax* liver stage parasites stain with rPvUIS4, we used this to establish a primary mask to define liver stage parasites, and then quantified the amount of EdA incorporated into each form using fluorescence intensity. This analysis revealed that, while most hypnozoite forms were negative for EdA, some forms of similar size and morphology were positive for EdA and were therefore synthesizing DNA at some point between days 5-8 dpi (Figure 3). We also assessed the effect of EdA and its cytotoxicity on PHH, which are non-replicative, and were not found to incorporate EdA. However, we noticed a slight cytotoxic effect of EdA on PHH count as compared to DMSO vehicle control. Cultures treated with 2 μM and 10 μM for three consecutive days had a slight statistically significant decrease in hepatocyte nuclei count, yet the toxic effect did not appear to hinder parasite growth (Figure 4c-d).

**Figure 4:**
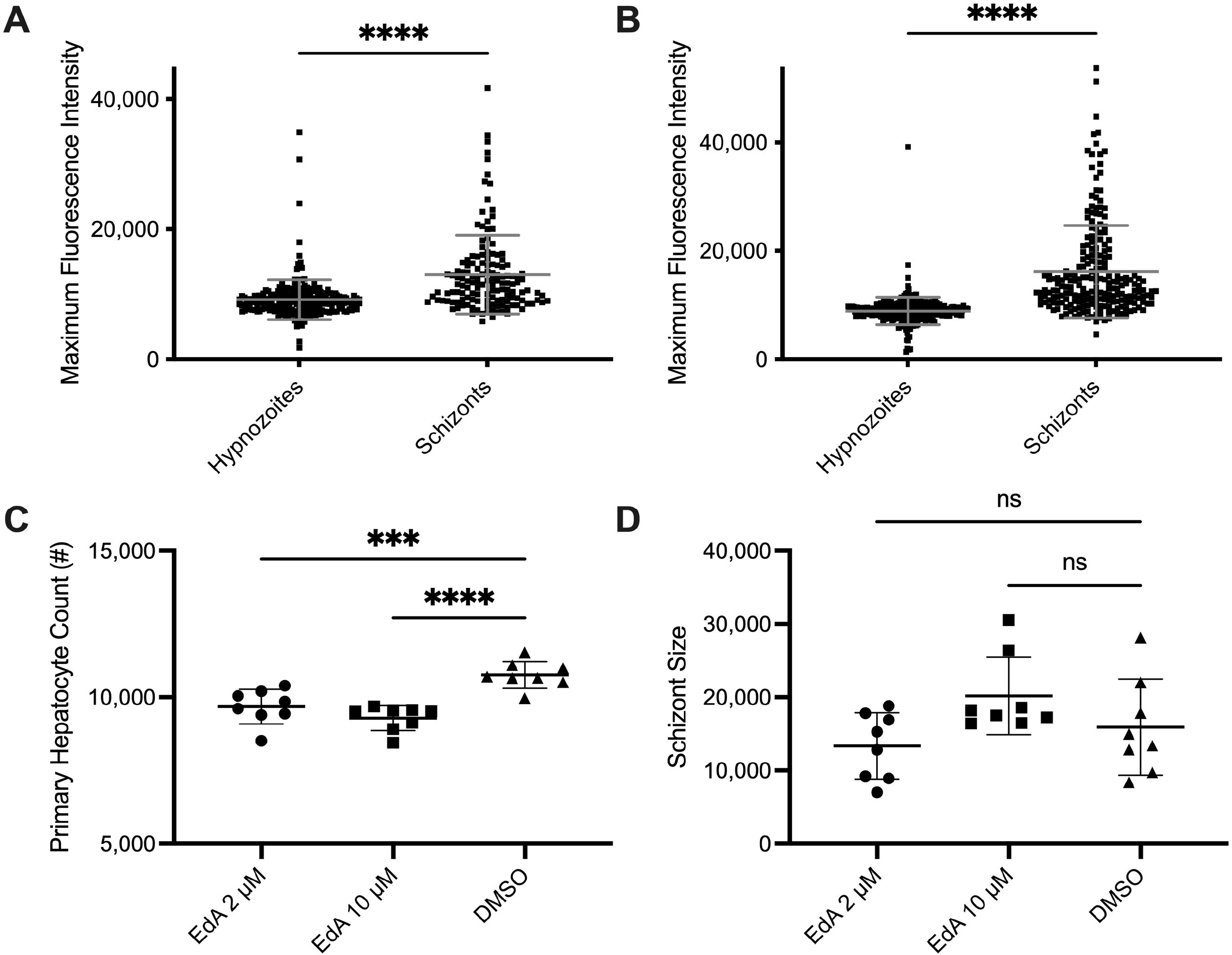
*P. vivax* hypnozoites and liver stage schizonts can be identified and quantified via EdA incorporation and high content imaging. Primary human hepatocytes (PHH) were seeded into a 384-well plate and infected with *P. vivax* sporozoites. **(A)** 2 μM or **(B)** 10 μM EdA was then supplemented on days 5, 6, 7, and 8 post infection. Parasites were stained with 1:25,000 PvUIS4, a parasitophorous vacuole membrane stain, and EdA was stained via a copper-catalyzed click reaction. PvUIS4 positive parasites were then separated into hypnozoites and liver stage schizonts by size using an ImageXpress confocal microscope. EdA incorporation was then assessed within each group using maximum fluorescence intensity. Parasites earlier deemed to be hypnozoites based on size but with high maximum fluorescence intensity for EdA were found to be early liver stage schizonts. **(C)** EdA cytotoxicity was assessed on PHH using nuclei count as an indicator of toxicity. **(D)** Liver stage schizont size was assessed for EdA vs. DMSO treated parasites. Figure is a representative of one independent replicate out of two (n = 2). Unpaired t-tests were used for statistical analysis in **A** and **B**. An ordinary one-way ANOVA Dunnett’s multiple comparisons test was used for **C**. *** p = 0.0005, **** p < 0.0001.

### 3.4 *P. berghei* liver stage schizonts incorporate EdA

*P. berghei* does not relapse *in vivo* and produces only liver stage schizonts. To confirm that all *P. berghei* liver stage parasites are actively replicating, and also confirm the incorporation of EdX in liver stages, we added EdA to *P. berghei* infected hepatocyte cultures for 24 hours prior to fixation. As shown in figure 5, all parasites were found to be positive for EdA staining. *P. berghei* liver stage assays are routinely performed by using the human hepatocarcinoma cell line (HepG2) as host cells [34]. Since HepG2 cells are actively replicating, we assessed HepG2 incorporation of EdA at 2 μM and 10 μM alone in the absence of infection for up to 72 hours (Supplemental Figure 1). We noted that while HepG2 cells successfully incorporate EdA, this modified purine is noticeably cytotoxic to the hepatocytes. While 60% of HepG2 cells incorporate EdA at the lower concentration of 2 μM, only 20% of HepG2 cells incorporate EdA at 10 μM (Supplemental Figure 2). HepG2 nuclei counts were significantly reduced at both concentrations as compared to DMSO control. Altogether, our results suggest that EdA can be used for shorter times at lower concentrations with HepG2 cells and can be used for longer durations with PHH.

**Figure 5:**
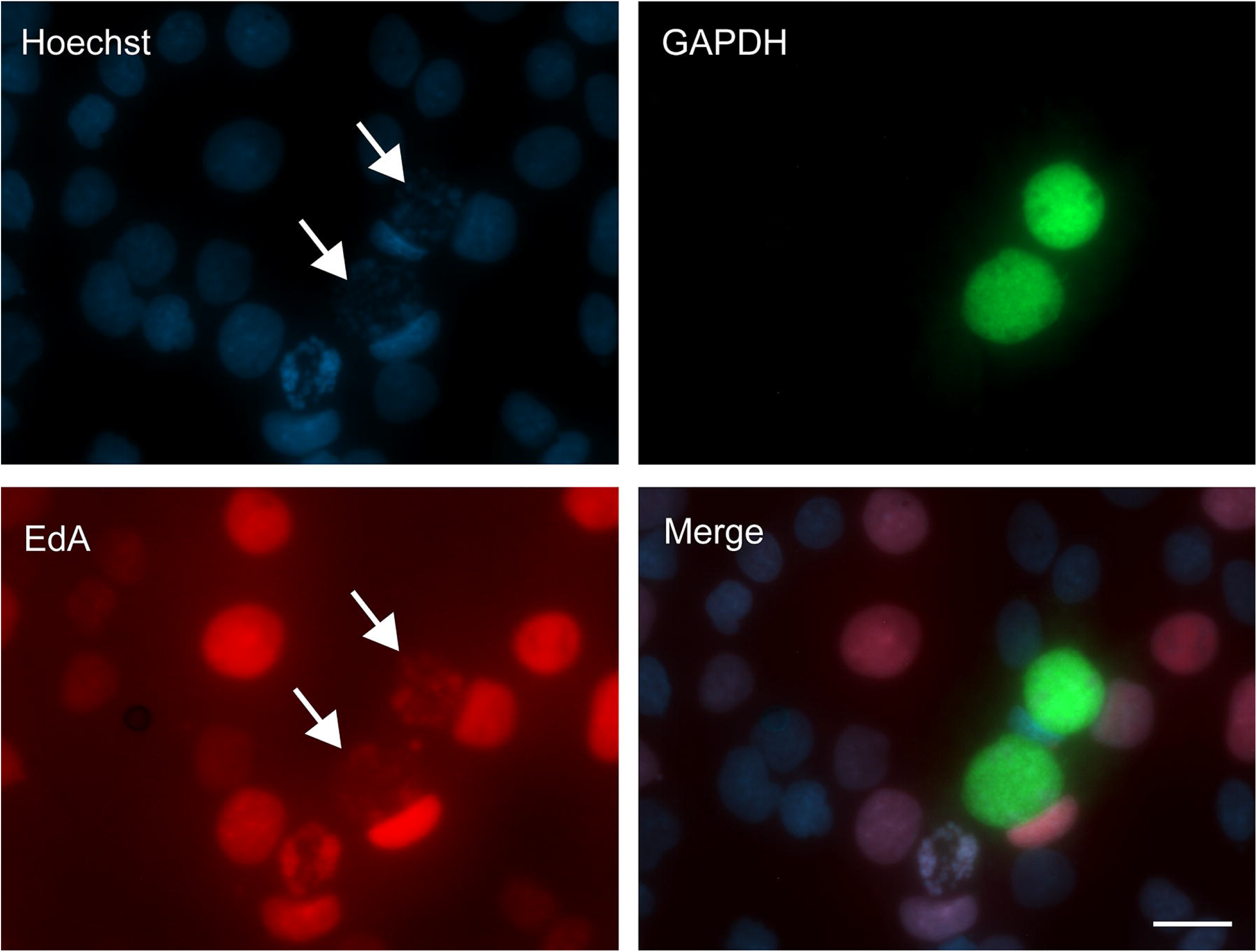
Alkyne modified adenosine (EdA) incorporates in replicating *P. berghei* liver stage parasites. HepG2 cells were infected with *P. berghei* sporozoites and 10μM EdA was supplemented at 24 hours post infection. Cells were then fixed 48 hours post infection. Detection of EdA was assessed via a copper-catalyzed click reaction (red). Parasites were co-stained with 1:10,000 GAPDH (green) and 10 μg/mL Hoechst 33342 (blue). Images were obtained on a Zeiss Axio Observer.Z1/7 microscope at 40x objective. Scalebar = 20 μm.

### 3.5 *P. falciparum* asexual blood stage parasites recrudescing from dihydroartemisinin (DHA) – induced dormancy do not incorporate EdA, but do incorporate ^[3H]^ hypoxanthine and ^[3H]^ adenosine

Previous studies have confirmed that exposure of ring stages to DHA induces a dormant phenotype that is both time of exposure and DHA concentration dependent [3, 5]. Due to the successful incorporation of EdA in *P. vivax* liver stage schizonts, we assessed if EdA incorporation could be used to differentiate DHA-induced dormant *P. falciparum* rings from recrudescing parasites. Following exposure to 700 nM DHA, parasites were sampled daily and pulsed with 10 μM EdA for 24 hours. We observed that as parasites recrudesced and the number of infected red blood cells (parasitemia) increased, incorporation of EdA did not increase concomitantly (Figure 6a). Thus, we assessed if this lack of correlation was due to potential purine storage in the parasite as a response to DHA treatment, or if DHA treated parasites became more sensitive to the alkyne modification to adenosine and thus were unable to uptake and incorporate EdA. Parasites were treated with DHA and then sampled daily and pulsed with either ^3^H-hypoxanthine or ^3^H-adenosine for 24 hours. Daily Giemsa smears were also collected to assess morphology of recrudescing parasites. To compare the incorporation of radiolabeled purines in recrudescing parasites, a control sample of parasites that did not undergo DHA treatment was included. The control sample of parasites received 24-hour pulses of radiolabeled hypoxanthine or adenosine. Control samples replicated and reached high parasitemia by day 2 and sampling was stopped. We observed that as parasites recrudesced and parasitemia increased, incorporation of radiolabeled purines occurred at the same rate as parasitemia measured by microscopic analysis of Giemsa-stained blood smears (Figure 6b). Altogether, our data suggest that either DHA exposed parasites do not store purines or that the alkyne modification affects incorporation into DNA of DHa treated parasites.

**Figure 6:**
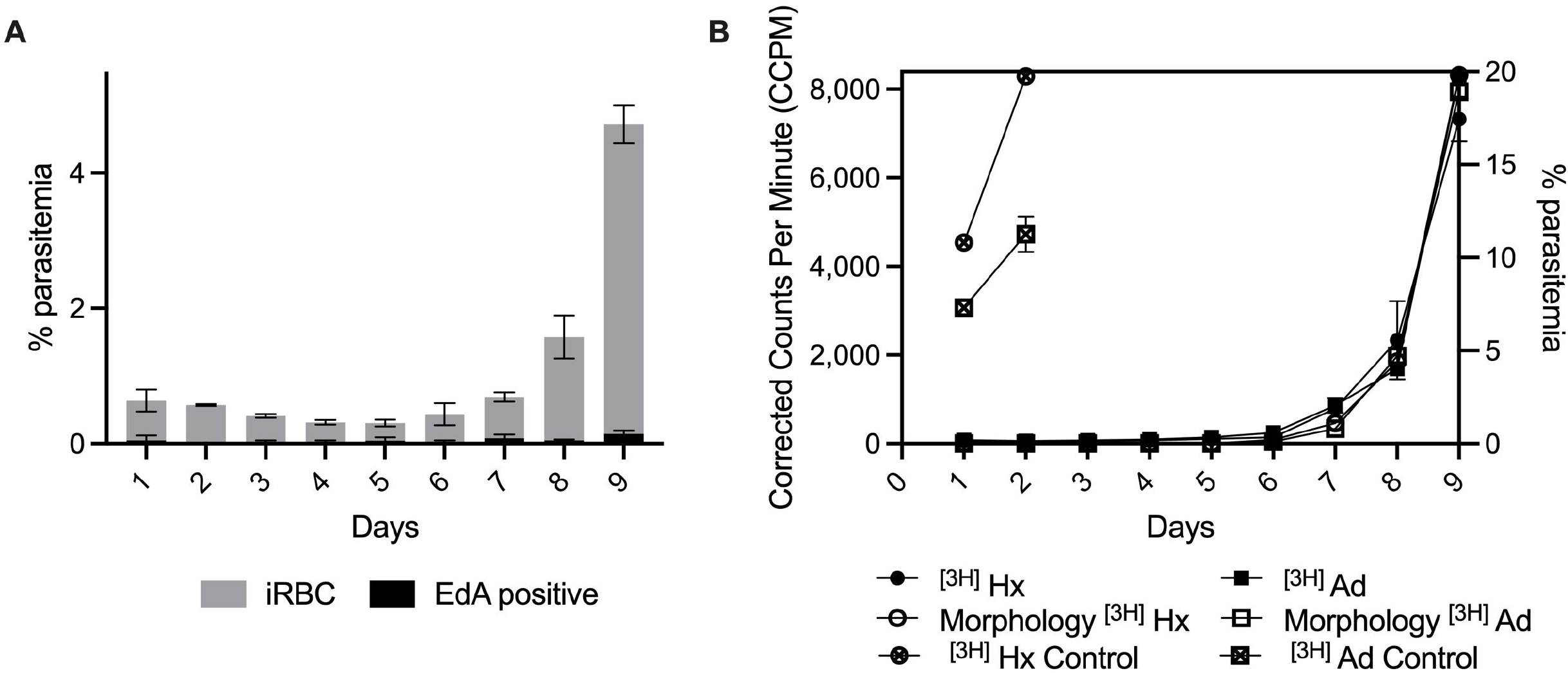
Asexual blood stage parasites recrudescing from DHA-induced dormancy incorporate [^3^H]-hypoxanthine and/or [^3^H]-adenosine but do not incorporate EdA. **A)** DHA dormancy was induced in *P. falciparum* asexual blood stage parasites and daily samples were acquired and pulsed with 10 μM EdA for 24 hours. Pulsed samples were analyzed via flow cytometry on a Beckman Coulter CytoFLEX. **B)** DHA dormancy was induced in *P. falciparum* asexual blood stage parasites. Daily samples were acquired and pulsed with [^3^H]-hypoxanthine (Hx) or [^3^H]-adenosine (Ad) for 24 hours. Daily Giemsa-stained smears were performed to assess recrudescence (% parasitemia). Control parasites were not treated with DHA. Radiolabeled samples were analyzed by a Perkin Elmer Microbeta^2^ scintillation counter. Date shown are one representative experiment of two independent experiments, (average ± SD).

## 4. DISCUSSION

The ability of *Plasmodium* to convert into a dormant phenotype and later reactivate causing recrudescent or relapse infections remains a serious barrier towards malaria eradication. Reactivation occurs both naturally in the liver stages and following a drug-induced growth arrest during the intraerythrocytic life cycle [35]. The mechanisms by which the parasite enters dormancy and later recrudesces or reactivates to continue development is not fully understood and we lack tools to study these mechanisms. For *P. vivax* liver stages, a H3K9Ac marker for nuclear division and a LISP2 marker have been developed and associated with reactivation [11, 12]. However, these markers are downstream of DNA synthesis and replication, and do not indicate the time at which exact DNA synthesis is initiated in activated hypnozoites. Since purine and pyrimidine nucleotides are the building blocks of nucleic acids, biosynthesis or incorporation of these building blocks marks initiation of DNA synthesis [36]. Since *Plasmodium* salvages purines from the host [26], we hypothesized that alkyne-containing purine nucleosides could be developed as makers for DNA synthesis markers to differentiate between active and dormant parasites. Our study provides the first report of a clickable DNA synthesis marker for *Plasmodium* which can easily be integrated into a staining panel design due to its flexibility. We show that EdA, EdI, 8eH, and EdH are incorporated into DNA in *P. falciparum* asexual blood stages, and that EdA is incorporated into actively replicating liver stage schizonts in *P. vivax* and *P. berghei,* but it does not incorporate into dormant *P. vivax* liver stage hypnozoites.

The methodology described in this study represents a valuable tool for *P. vivax* liver stage studies as it is the first to describe a DNA synthesis marker which can be used to distinguish actively replicating and dormant liver stages of the parasite. Interestingly, we noted that some liver forms of similar size and morphology to that of PI4K-insensitive hypnozoites do incorporate DNA, indicating that these forms could be newly reactivating parasites (Figure 3). Alternatively, recent reports describe how liver stage parasites must constantly buffer themselves against host cell lysosomes [37]. Thus, it is also possible that these small, EdA-positive liver stage forms are schizonts which failed to develop due to factors such as the host response to infection. Our data suggest EdA could be used in future studies to identify and characterize newly reactivating parasites and other host-parasite interactions. Interestingly, only EdA was found to incorporate in *P. vivax* and *P. berghei* liver stage parasites. Previous work has shown rapid metabolism of inosine and hypoxanthine into allantoin by rat hepatocytes, which could explain why parasite incorporation of EdI, 8eH, or EdH was not achieved in the liver stage [38].

On the other hand, we observed that active blood stage *P. falciparum* parasites incorporated all the alkyne-modified purines (Figure 1). This creates a novel opportunity to study DNA synthesis pathways and cell cycling. Unexpectedly, *P. falciparum* recrudescing blood stage parasites coming out of DHA-induced dormancy did not incorporate any of the modified purines (Figure 6). Moreover, when comparing incorporation of radiolabeled hypoxanthine and adenosine, to EdA incorporation in recrudescing parasites, we found that EdA was not incorporated while the radiolabeled purines were metabolized and incorporated into DNA. Previous work has shown that artemisinin-resistant *P. falciparum* has altered metabolic programming [39]. Blood stage parasites showed decreased levels of adenosine deaminase and hypoxanthine-guanine phosphoribosyltransferase (HGXPRT) in Kelch13 (K13) mutant parasites, suggesting a prior pooling of purines in response to ART drug exposure [39]. While the work reported here was conducted utilizing a K13-wild type *P. falciparum* strain sensitive to ART, it is possible that in response to DHA exposure, even K13-wild type parasites may accumulate internal stores of purines after exposure to DHA. Alternatively, structural modifications to purine precursors may cause steric hinderance of the purine salvage enzymes (e.g., purine nucleoside phosphorylase). Although the EdX may not be useful for DHA-induced dormancy studies in *P. falciparum,* they remain valuable as an alternative to radiolabeled purines, as these alkyne-modified purines can potentially be used as an inexpensive tool for investigating the parasite’s scavenging mechanisms.

In *P. falciparum* blood stage *in vitro* cultures, we observed that in the presence of hypoxanthine, the modified purines were not cytotoxic. However, parasites lacking supplementation of hypoxanthine for purine salvage were not able to sustain growth when supplemented with the alkyne modified purines alone. Nevertheless, incorporation of EdA, EdI, 8eH, and EdH was observed in media supplemented with hypoxanthine. These data suggest that parasite DNA replication enzymes sense a difference between modified and unmodified purines; however, incorporation still occurs at a sufficient low enough rate that it cannot replace hypoxanthine to sustain replication and parasite growth, but it can be visualized via fluorescence microscopy. In our *P. vivax* and *P. berghei* liver stage experiments, it is important to note that a slight hepatocyte toxicity from EdA was observed which could confound our studies. Nevertheless, parasite growth remained unaffected (figure 4). In the present study, long incubations times such as 48-72 hours were used. However, to assess reactivation from dormancy, shorter incubation times could be used in future studies to obtain narrower time windows of reactivation and to limit negative impacts on the assay due to hepatocyte toxicity.

This methodology provides many opportunities for the study of malaria parasite biology. While most preliminary *in vitro* drug screening for liver stage activity is conducted using *P. berghei,* medium throughput platforms using high-content imaging of *P. vivax* liver stages are now being used to confirm and optimize anti-hypnozoite hits [8]. By incorporating EdA into the high-content pipeline, we can further characterize both the parasite forms quantified during analysis, as well as gain a better understanding of the effect of agonists or antagonists of hypnozoite reactivation. Ideally, future experimentation will involve a *P. vivax* time course with combination of LISP2, EdA, and H3K9Ac staining to better define the timing of DNA replication, nuclear division, and membrane synthesis in reactivating parasites. Furthermore, EdA staining approach could aid in elucidating the cause of hypnozoite reactivation, which has been hypothesized to include fever, hemolysis, malaria reinfection, and chemical reactivation [40, 41]. Recently we reported the recruitment of host aquaporin 3 to the parasite PVM, as well as the formation of a tubulovesicular network around *P. vivax* liver forms; these mechanisms have been hypothesized to be part of the parasite’s nutrient-scavenging pathways [10, 42]. EdA staining could be used as a bait to further elucidate pathways responsible for purine scavenging. Additionally, we can postulate that EdA would incorporate into *P. cynomolgi* liver stages and could therefore be useful for *P. cynomolgi* drug discovery platforms. *P. cynomolgi* produces hypnozoites, although *P. cynomolgi* hypnozoites and liver stage schizonts have been reported to be much smaller than their *P. vivax* counterparts [43–45]. Yet, size alone is often use as the defining attribute of hypnozoites versus schizonts during high-content analysis. Given our finding that some hypnozoite-like forms are synthesizing DNA, and *P. cynomolgi* liver forms are both relatively smaller than those of *P. vivax,* misclassification of hypnozoites and schizonts is very possible in this model as well, but it could be better characterized using a marker such as EdA.

Herein, we detail the synthesis of new alkyne-modified purines, including inosine (EdI) and hypoxanthine (EdH and 8eH) derivatives, and the first reported use of alkyne modified purines to study the biology of *Plasmodium.* The use of alkyne-modified purines enables Chemo-labelling (“Click chemistry”), providing advantages over traditional antibody staining in that researchers do not have to consider animal/species cross-reactivity. Furthermore, it offers increased flexibility since the alkyne modified purines can be “clicked” to any azide-linked molecule of choice and can also be leveraged in a high-throughput manner using high-content imaging systems. In addition, the novel purine analogues reported here may have potential uses in other organisms that have yet to be tested and validated. This new tool is inexpensive, easy to incorporate into current workflows, and provides flexibility, making it an ideal tool as a DNA synthesis marker.

## ACKNOWLEDGEMENTS

We would like to thank Dr. Muthugapatti Kandasamy and the University of Georgia Biomedical Microscopy Core for training and use of the DeltaVision II deconvolution microscope and ImageXpress Micro Confocal system, provided with support from the Georgia Research Alliance. We would also like to thank Julie Nelson and the Center for Tropical and Emerging Global Diseases Cytometry Shared Resource Laboratory for the training and use of the Beckman Coulter CytoFLEX. We thank the malaria patients of northeastern Cambodia for participation in this study. We thank Dr. Ashutosh K. Pathak and the University of Georgia Sporocore team as well as the insectary team at the Institut Pasteur of Cambodia for the production of *P. berghei* and *P. vivax* sporozoite-infected mosquitos, respectively.

## AUTHOR CONTRIBUTIONS

Conceptualization, AB, GL, JL and DK. Methodology, AB, GL, SM, and JS. Formal Analysis, AB, GL, JL, and DK. Design and synthesis of purine analogs, GL, DD, CY, and JL. Resources, AV, BW, EFM, MBC, JL, and DK. Writing – Original Draft, AB and GL. Writing – Review and Editing, AB, GL, SM, AV, BW, JS, EFM, MBC, JL, and DK. Visualization, AB and JL. Supervision, JL and DK. Funding Acquisition, JL and DK. All authors contributed to the article and approved the submitted version.

## FUNDING AND ADDITIONAL INFORMATION

Funding support was provided by the Bill & Melinda Gates Foundation (OPP1023601 to DEK), the Georgia Research Alliance (DEK), and fellowship support by the NIH (T32AI060546, to AB).

## CONFLICT OF INTEREST

The authors declare no competing interests.

---

**Supplemental Figure 1: Alkyne modified adenosine (EdA) incorporates in replicating HepG2 mammalian cells.** HepG2 cells were seeded at 5,000 cells/well and 2 μM or 10 μM EdA was then supplemented at 24 hours post-seed. Cells were then fixed 72 hours post-seed. Detection of EdA was assessed via a copper-catalyzed click reaction (green). HepG2 nuclei were co-stained with 10 μg/mL Hoechst 33342 (blue). Images were obtained on a Lionheart FX automated microscope at 10x objective.

**Supplemental Figure 2: EdA labels replicating HepG2 mammalian cells and is cytotoxic.** HepG2 cells were seeded at 5,000 cells/well and 2 μM or 10 μM EdA was supplemented at 24 hours post-seed. Cells were fixed 72 hours post-seed. EdA incorporation was assessed via a copper-catalyzed click reaction and HepG2 nuclei were co-stained with 10 μg/mL Hoechst 33342. Analysis was then conducted using Gen5 software. **A**) Nuclei count was assessed by Hoechst nuclear staining and **B**) EdA incorporation was assessed as percentage of EdA positive nuclei. Data shown are one representative experiment of two independent experiments (average ± SD). Significance was assessed using an ordinary one-way ANOVA with Dunnett’s multiple comparisons test (**A**) or an unpaired t-test (**B**), **** p < 0.0001.

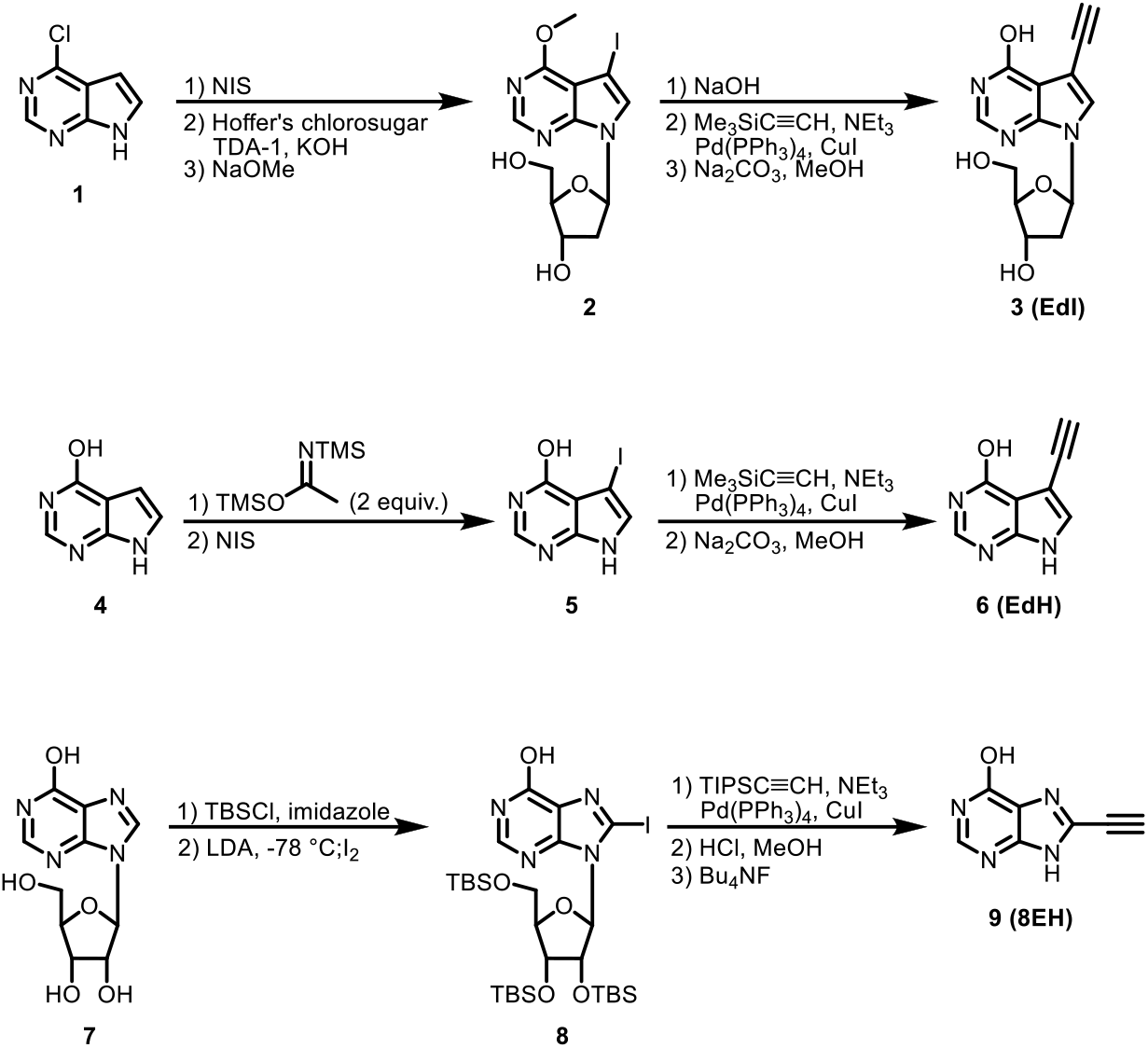

## Notes

### Competing Interest Statement

The authors have declared no competing interest.

## REFERENCES

1. World Malaria Report 2020. World Health Organization, 2020.

2. Krotoski, W.A., et al., Demonstration of hypnozoites in sporozoite-transmitted Plasmodium vivax infection. Am J Trop Med Hyg, 1982. 31(6): p. 1291–3.

3. Teuscher, F., et al., Phenotypic changes in artemisinin-resistant Plasmodium falciparum lines in vitro: evidence for decreased sensitivity to dormancy and growth inhibition. Antimicrob Agents Chemother, 2012. 56(1): p. 428–31.

4. Witkowski, B., et al., Increased tolerance to artemisinin in Plasmodium falciparum is mediated by a quiescence mechanism. Antimicrob Agents Chemother, 2010. 54(5): p. 1872–7.

5. Tucker, M.S., et al., Phenotypic and genotypic analysis of in vitro-selected artemisinin-resistant progeny of Plasmodium falciparum. Antimicrob Agents Chemother, 2012. 56(1): p. 302–14.

6. Baird, J.K., Primaquine toxicity forestalls effective therapeutic management of the endemic malarias. Int J Parasitol, 2012. 42(12): p. 1049–54.

7. Baird, J.K., Tafenoquine for travelers’ malaria: evidence, rationale and recommendations. J Travel Med, 2018. 25(1).

8. Roth, A., et al., A comprehensive model for assessment of liver stage therapies targeting Plasmodium vivax and Plasmodium falciparum. Nature communications, 2018. 9(1): p. 1837–1837.

9. Schafer, C., et al., A recombinant antibody against Plasmodium vivax UIS4 for distinguishing replicating from dormant liver stages. Malaria journal, 2018. 17(1): p. 370–370.

10. Sylvester, K., et al., Characterization of the Tubovesicular Network in Plasmodium vivax Liver Stage Hypnozoites and Schizonts. Front Cell Infect Microbiol, 2021. 11: p. 687019.

11. Gupta, D.K., et al., The Plasmodium liver-specific protein 2 (LISP2) is an early marker of liver stage development. eLife, 2019. 8: p. e43362.

12. Mikolajczak, S.A., et al., Plasmodium vivax liver stage development and hypnozoite persistence in human liver-chimeric mice. Cell host & microbe, 2015. 17(4): p. 526–535.

13. Voorberg-van der Wel, A.M., et al., A dual fluorescent Plasmodium cynomolgi reporter line reveals in vitro malaria hypnozoite reactivation. Commun Biol, 2020. 3: p. 7.

14. WHO Guidelines for malaria. Geneva: World Health Organization.

15. Dondorp, A.M., et al., Artemisinin resistance in Plasmodium falciparum malaria. N Engl J Med, 2009. 361(5): p. 455–67.

16. Phyo, A.P., et al., Declining Efficacy of Artemisinin Combination Therapy Against P. Falciparum Malaria on the Thai-Myanmar Border (2003-2013): The Role of Parasite Genetic Factors. Clinical infectious diseases: an official publication of the Infectious Diseases Society of America, 2016. 63(6): p. 784–791.

17. Leang, R., et al., Evidence of Plasmodium falciparum Malaria Multidrug Resistance to Artemisinin and Piperaquine in Western Cambodia: Dihydroartemisinin-Piperaquine Open-Label Multicenter Clinical Assessment. Antimicrobial agents and chemotherapy, 2015. 59(8): p. 4719–4726.

18. Phyo, A.P., et al., Emergence of artemisinin-resistant malaria on the western border of Thailand: a longitudinal study. Lancet (London, England), 2012. 379(9830): p. 1960–1966.

19. Teuscher, F., et al., Artemisinin induced dormancy in plasmodium falciparum: duration, recovery rates, and implications in treatment failure. J Infect Dis, 2010. 202(9): p. 1362–8.

20. Codd, A., et al., Artemisinin-induced parasite dormancy: a plausible mechanism for treatment failure. Malaria journal, 2011. 10: p. 56–56.

21. Assefa, D.G., G. Yismaw, and E. Makonnen, Efficacy of dihydroartemisinin-piperaquine versus artemether-lumefantrine for the treatment of uncomplicated Plasmodium falciparum malaria among children in Africa: a systematic review and meta-analysis of randomized control trials. Malaria journal, 2021. 20(1): p. 340–340.

22. Gratzner, H.G., Monoclonal antibody to 5-bromo- and 5-iododeoxyuridine: A new reagent for detection of DNA replication. Science, 1982. 218(4571): p. 474–5.

23. Salic, A. and T.J. Mitchison, A chemical method for fast and sensitive detection of DNA synthesis in vivo. Proc Natl Acad Sci U S A, 2008. 105(7): p. 2415–20.

24. Janse, C.J., et al., Bromo-deoxyuridine is not incorporated into DNA of malaria parasites. Trans R Soc Trop Med Hyg, 1991. 85(6): p. 727–8.

25. Rager, N., et al., Localization of the Plasmodium falciparum PfNT1 nucleoside transporter to the parasite plasma membrane. J Biol Chem, 2001. 276(44): p. 41095–9.

26. Downie, M.J., K. Kirk, and C.B. Mamoun, Purine Salvage Pathways in the Intraerythrocytic Malaria Parasite Plasmodium falciparum. Eukaryotic Cell, 2008. 7(8): p. 1231–1237.

27. Merrick, C.J., Transfection with thymidine kinase permits bromodeoxyuridine labelling of DNA replication in the human malaria parasite Plasmodium falciparum. Malar J, 2015. 14: p. 490.

28. Cassera, M.B., et al., Purine and pyrimidine pathways as targets in Plasmodium falciparum. Curr Top Med Chem, 2011. 11(16): p. 2103–15.

29. Neef, A.B., F. Samain, and N.W. Luedtke, Metabolic Labeling of DNA by Purine Analogues in Vivo. ChemBioChem, 2012. 13(12): p. 1750–1753.

30. Pawlowic, M.C., et al., Genetic ablation of purine salvage in Cryptosporidium parvum reveals nucleotide uptake from the host cell. Proc Natl Acad Sci U S A, 2019. 116(42): p. 21160–21165.

31. Trager, W. and J.B. Jensen, Human malaria parasites in continuous culture. Science, 1976. 193(4254): p. 673–5.

32. Balu, B., et al., A genetic screen for attenuated growth identifies genes crucial for intraerythrocytic development of Plasmodium falciparum. PLoS One, 2010. 5(10): p. e13282.

33. Mueller, I., et al., Key gaps in the knowledge of Plasmodium vivax, a neglected human malaria parasite. The Lancet Infectious Diseases, 2009. 9(9): p. 555–566.

34. Dorjsuren, D., et al., Chemoprotective antimalarials identified through quantitative high-throughput screening of Plasmodium blood and liver stage parasites. Scientific reports, 2021. 11(1): p. 2121–2121.

35. Barrett, M.P., et al., Protozoan persister-like cells and drug treatment failure. Nat Rev Microbiol, 2019. 17(10): p. 607–620.

36. Hyde, J.E., Targeting purine and pyrimidine metabolism in human apicomplexan parasites. Current drug targets, 2007. 8(1): p. 31–47.

37. Niklaus, L., et al., Deciphering host lysosome-mediated elimination of Plasmodium berghei liver stage parasites. Scientific Reports, 2019. 9(1): p. 7967.

38. Vincent, M.F., G. Van den Berghe, and H.G. Hers, Metabolism of hypoxanthine in isolated rat hepatocytes. The Biochemical journal, 1984. 222(1): p. 145–155.

39. Mok, S., et al., Artemisinin-resistant K13 mutations rewire Plasmodium falciparum’s intra-erythrocytic metabolic program to enhance survival. Nature communications, 2021. 12(1): p. 530–530.

40. Adekunle, A.I., et al., Modeling the dynamics of Plasmodium vivax infection and hypnozoite reactivation in vivo. PLoS neglected tropical diseases, 2015. 9(3): p. e0003595–e0003595.

41. Shanks, G. and N. White, The activation of vivax malaria hypnozoites by infectious diseases. The Lancet infectious diseases, 2013. 13.

42. Posfai, D., et al., Plasmodium vivax Liver and Blood Stages Recruit the Druggable Host Membrane Channel Aquaporin-3. Cell Chem Biol, 2020. 27(6): p. 719–727.e5.

43. Dembele, L., et al., Towards an In Vitro Model of Plasmodium Hypnozoites Suitable for Drug Discovery. PLOS ONE, 2011. 6(3): p. e18162.

44. Dembélé, L., et al., Persistence and activation of malaria hypnozoites in long-term primary hepatocyte cultures. Nature Medicine, 2014. 20(3): p. 307–312.

45. Voorberg-van der Wel, A., et al., Transgenic Fluorescent Plasmodium cynomolgi Liver Stages Enable Live Imaging and Purification of Malaria Hypnozoite-Forms. PLOS ONE, 2013. 8(1): p. e54888.

