## Supplemental Figures for "Alkyne modified purines for assessing activation of *Plasmodium vivax* hypnozoites and growth of pre-erythrocytic and erythrocytic stages in *Plasmodium* spp"

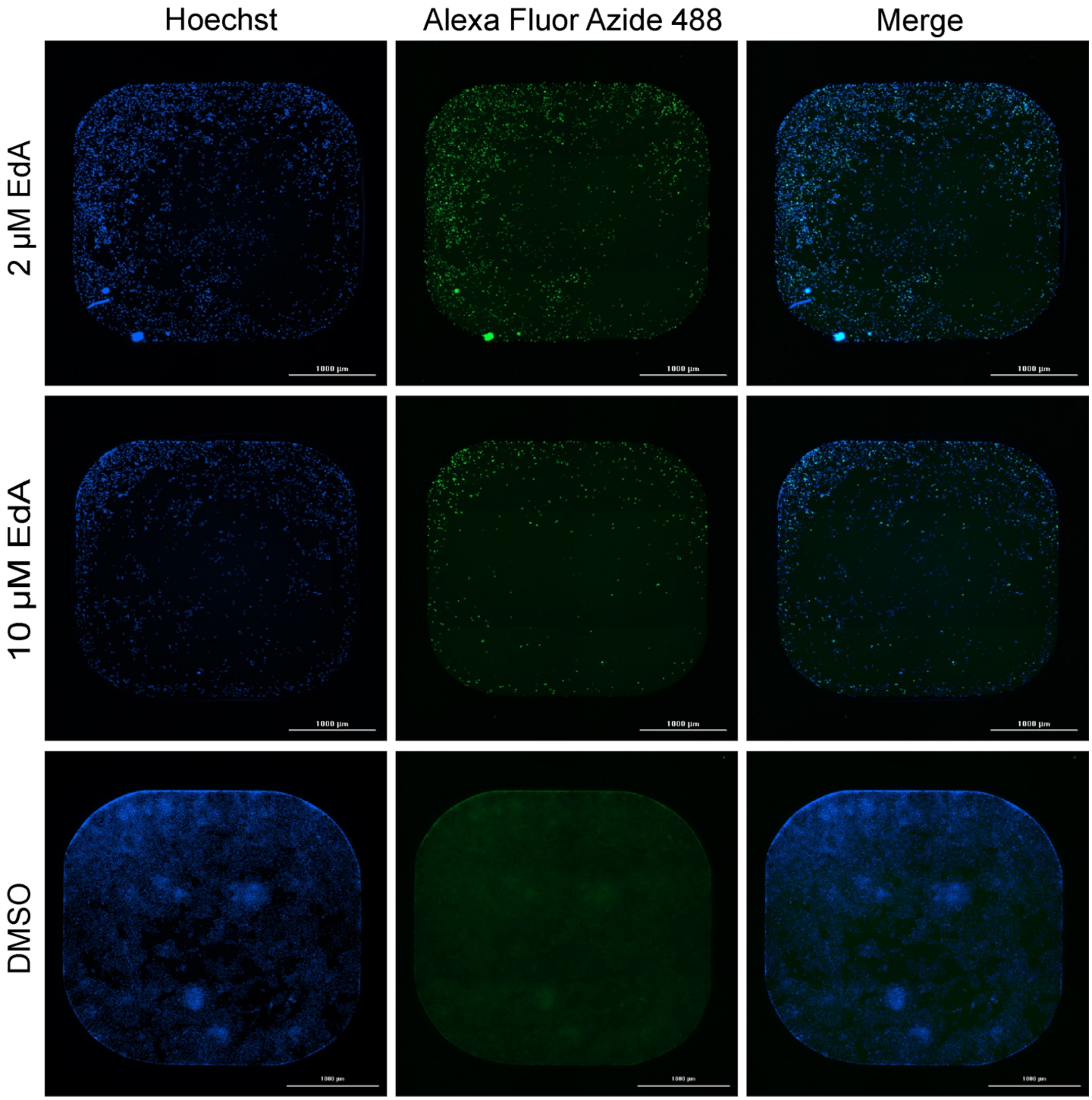


**Supplemental Figure 1: Alkyne modified adenosine (EdA) incorporates in replicating HepG2 mammalian cells**. HepG2 cells were seeded at 5,000 cells/well and 2 µM or 10 µM EdA was then supplemented at 24 hours post-seed. Cells were then fixed 72 hours post-seed. Detection of EdA was assessed via a copper-catalyzed click reaction (green). HepG2 nuclei were co-stained with 10 µg/mL Hoechst 33342 (blue). Images were obtained on a Lionheart FX automated microscope at 10x objective.

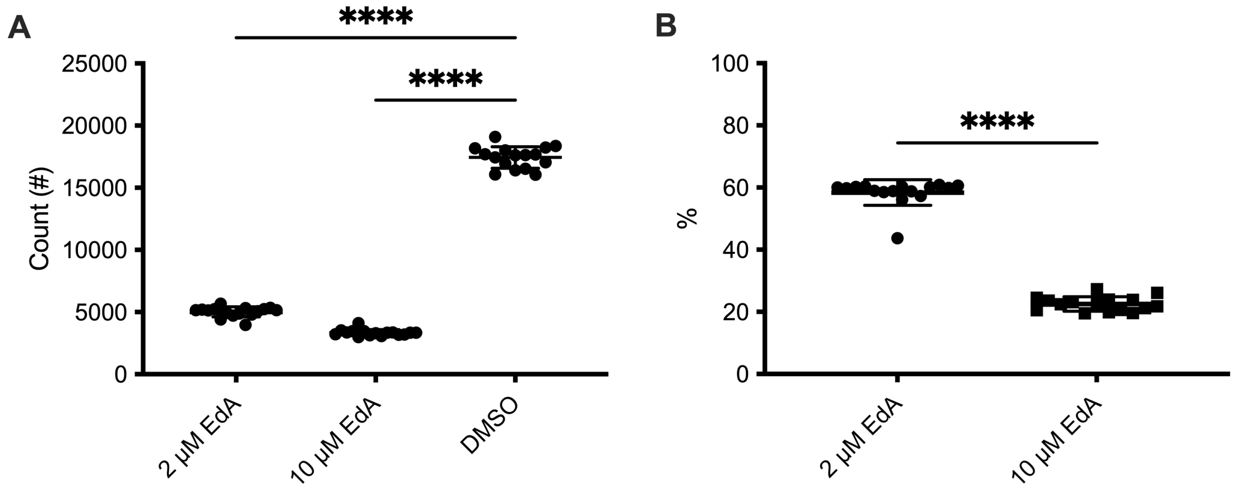


**Supplemental Figure 2: EdA labels replicating HepG2 mammalian cells and is cytotoxic.** HepG2 cells were seeded at 5,000 cells/well and 2 µM or 10 µM EdA was supplemented at 24 hours post-seed. Cells were fixed 72 hours post-seed. EdA incorporation was assessed via a copper-catalyzed click reaction and HepG2 nuclei were co-stained with 10 µg/mL Hoechst 33342. Analysis was then conducted using Gen5 software. **A)** Nuclei count was assessed by Hoechst nuclear staining and **B)** EdA incorporation was assessed as percentage of EdA positive nuclei. Data shown are one representative experiment of two independent experiments (average ± SD). Significance was assessed using an ordinary one-way ANOVA with Dunnett's multiple comparisons test (**A)** or an unpaired t-test (**B**), **** $p < 0.0001$.
